# The three regimes of spatial recovery

**DOI:** 10.1101/308635

**Authors:** Yuval R. Zelnik, Jean-Francois Arnoldi, Michel Loreau

## Abstract

An enduring challenge for ecology is identifying the drivers of ecosystem and population stability. In a spatially explicit context, key features to consider are landscape spatial structure, local interactions, and dispersal. Substantial work has been done on each of these features as a driver of stability, but little is known on the interplay between them. Missing has been a more integrative approach, able to map and identify different dynamical regimes, and thus to predict a system’s response to perturbations. Here we first consider a simple scenario, i.e. the recovery of a homogeneous metapopulation from a single localized pulse disturbance. The analysis of this scenario reveals three fundamental recovery regimes: *Isolated Regime* when dispersal is not significant, *Rescue Regime* when dispersal mediates recovery, and *Mixing Regime* when perturbations spread throughout the system. Despite its simplicity, our approach leads to remarkably general predictions. These include the qualitatively different outcomes of various scenarios of habitat fragmentation, the surprising benefits of local extinctions on population persistence at the transition between regimes, and the productivity shifts of metacommunities in a changing environment. This study thus provides context to known results and insight into future directions of research.

## I. INTRODUCTION

How can dispersal, ecosystem size and local dynamics interact to determine recovery from a disturbance? This question is fundamental to ecology not only due to its relevance for conservation and management, but because it connects key concepts of ecology, such as stability, landscape, metapopulations, and disturbance.

Dispersal plays a fundamental role in all aspects of ecology, affecting the stability of populations [1], biodiversity patterns [18], trophic interactions [29] and evolutionary dynamics [4]. Dispersal is most often studied because of two effects it has on ecosystems: sustaining diversity [20] and generating population synchrony [1, 21]. When dispersal is weak it can promote diversity, allowing populations to benefit from spatial insurance effects, whereby good patches prevent local extinctions in less favored locations [25]. This effect is fundamental in the context of biodiversity loss caused by human-induced landscape fragmentation, which impedes dispersal [7, 11]. Dispersal, however, is not always beneficial. Strong dispersal may synchronize population dynamics and cause global extinctions. It can inhibit spatial insurance effects, causing generalist species to competitively exclude specialists [1]. These two cases are extremes where there is a clear separation of timescales between the local dynamics and the time it takes to disperse across the system. In between is an intermediate regime without a clear separation of timescales, which has not been investigated much or even well defined.

Not all relevant spatial aspects of ecosystems are centered on dispersal and interactions across space. Sheer size is also important as spatial processes are effectively mediated by the system size [14]. Larger regions can allow for substantial spatial heterogeneity, from asynchrony due to nonlinear local dynamics or disturbance regimes [6, 16], to an imposed structure due to topography or climatic gradients [31]. Spatial averaging over such heterogeneities has lead to many well-known concepts in ecology, such as the species-area relationship [8] and landscape equilibrium [36].

The ecological concepts discussed above, such as synchrony and spatial averaging, are non-trivial due to local dynamics that act in conjunction with spatial effects. In fact, ecology has long focused on local non-spatial behavior, with central issues such the diversity-stability debate [24, 26–28] largely addressed by focusing on local interactions between species. Assumptions on local dynamics vary greatly from linear behavior around an equilibrium to highly nonlinear dynamics far from it. This is evident in stability research where noisy time-series are typically assumed to be close to equilibrium [10], while catastrophic regime shifts are inherently nonlinear [34]. When considering spatial aspects, however, most studies implicitly or explicitly assume linear behavior, while research into nonlinear behavior is mostly focused on specific scenarios such as emergent stationary spatial patterns in drylands [37] or chaotic behavior of algae blooms [13].

Stability is a central notion in ecology, yet its very definition is strongly debated, as is the preferred method of measuring it in both theoretical and empirical work [10]. Nevertheless, the concept of stability comes down fundamentally to the ability of the system to recover from a perturbation [2], which may be affected by the timing of the perturbation (e.g. constant or single event), the dynamical aspect considered (e.g. rate of convergence or disturbance strength withstood), or the central measure recovered (e.g. biodiversity or overall biomass).

Recent work has investigated the stability of populations and ecosystems in a spatial context, by explicitly considering the issues of dispersal, system size and local dynamics [9, 12, 15, 30, 38]. The scenarios considered, however, are often quite specific, and it is difficult do draw general conclusions from them. Moreover, since there is no clear framework in which to understand the phenomena described, results are hard to compare, limiting the potential for synthesis. Here we propose that answering the preliminary question of how dispersal, ecosystem size and local dynamics interact to determine recovery from a disturbance provides a unifying perspective on the various regimes of recovery.

We first address this question in a simplified setting, i.e. the response of a single species at equilibrium to a disturbance (a sudden change in abundance) in a uniform one-dimensional landscape. We monitor the time needed for the system to return to its pre-disturbed state. This notion is easily measured and understood, while having clear relations to other notions of stability [3]. This allows us to draw an exhaustive map of dynamical behavior, predicting the transitions between three qualitatively different recovery regimes: *Isolated Regime* (IR), *Rescue Regime* (RR) and *Mixing Regime* (MR). In IR dispersal is not essential for recovery, in RR propagation of biomass is key for recovery, and in MR recovery occurs after the disturbance’s effect has spread throughout the system.

We then translate this approach to more complex ecological settings: early warning signals of catastrophic transitions, the interplay between local extinctions and metapopulation persistence, and productivity of metacommunities in a changing environment. Our approach thus defines a powerful methodology for the analysis of spatial ecosystems, and proposes new directions of research for basic theory, field studies and conservation.

## II. CONCEPTUAL FRAMEWORK

In order to understand the spatial nature of stability, we focus on the behavior of a single species that is at a stable equilibrium before a disturbance occurs. We use a partial differential equation as a mathematical description of our system, so that its behavior is governed by the combination of local interactions of individuals, and dispersal in space of these individuals. The dynamics of the system read:

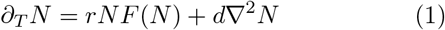

with *N* the density of individuals at a location, *r* the local dynamics rate, and *d* the dispersal coefficient. The time derivative is *∂_T_*, the diffusion operator is ∇^2^ (in a one dimensional system it amounts to the second spatial derivative *∂_XX_*), and *F* is some (typically nonlinear) function that describes the overall effect of local interactions between individuals, which we assume has an equilibrium solution for *N* = *K*. If for example *F(N)* = (1 — *N/K*) with *K* the carrying capacity, then the system’s dynamics is logistic growth. We denote the size of the system by *L*.

An important feature of the system is the nonlinearity of its local dynamics, and in particular how much slower are local dynamics when far from equilibrium.

Concretely we focus on one specific type of local dynamics given by *F(N)* = (1 — *N/K*)(*N/K*)^*γ*^, where large *γ* leads to higher nonlinearity and slower dynamics far from equilibrium *N* = *K*.

Having defined the system, we now study its recovery from a single, localized, pulse disturbance. In Fig. 1 we show the different possible responses of the system, and how these map out in the parameter space of *r* and *d*. We find that the recovery typically follows one of the three scenarios: Isolated Regime (IR), Rescue Regime (RR), and Mixing Regime (MR).

For weak dispersal *d* and fast local dynamics *r*, the system is in IR, and the recovery of each location occurs without any relation to other regions. We quantify this notion by the ratio between the biomass recovery when *d* = 0 and when *d* > 0, within the same time frame. On the other hand, for large *d* and low *r* values, the system is in MR, and the recovery of all the locations of the system effectively occurs together: dispersal homogenizes the system before local recovery processes take place. This can be quantified by defining a mixing time, when all locations of the system reach similar biomass densities (within some threshold), and attributing all biomass recovered after this time to MR. Finally, for intermediate values of *d* and *r*, the system can be in RR, where undisturbed regions aid the disturbed ones via spatial spread processes. This occurs when non-spatial dynamics of the disturbed region are sufficiently slow, so that spatial recovery processes (e.g. front propagation) can come into play, yet dispersal is not strong enough to homogenize the system. While it is possible to define this recovery directly, we find it simpler to define it as any recovery that is not attributed to IR or MR.

An important determinant of recovery is the disturbance itself. Here there are essentially three disturbance properties to consider: overall strength *s* (total biomass removed), spatial extent *σ* (size of disturbed region), and intensity *ρ* = *s/σ* (biomass removed per area in the disturbed region). As shown in Fig. 2, These properties define a parameter space in which we can map all possible disturbances, and the system response they induce. For a given disturbance strength *s* there are two extreme cases (orange circles in Fig. 2a): a global disturbance where the extent *σ* is maximal (the entire system is disturbed), and a localized disturbance where the intensity *ρ* is maximal (all biomass is removed from the disturbed region). These contrasting cases are met with very different recovery behaviors. For example, for a given disturbance strength the recovery timescale is much shorter in response to a global disturbance than to a localized one. Moreover, the transient spatial profiles are strikingly different: in Fig. 2b we see the formation of fronts, while Fig. 2d only shows a uniform increase of biomass in time.

**FIG. 1:**
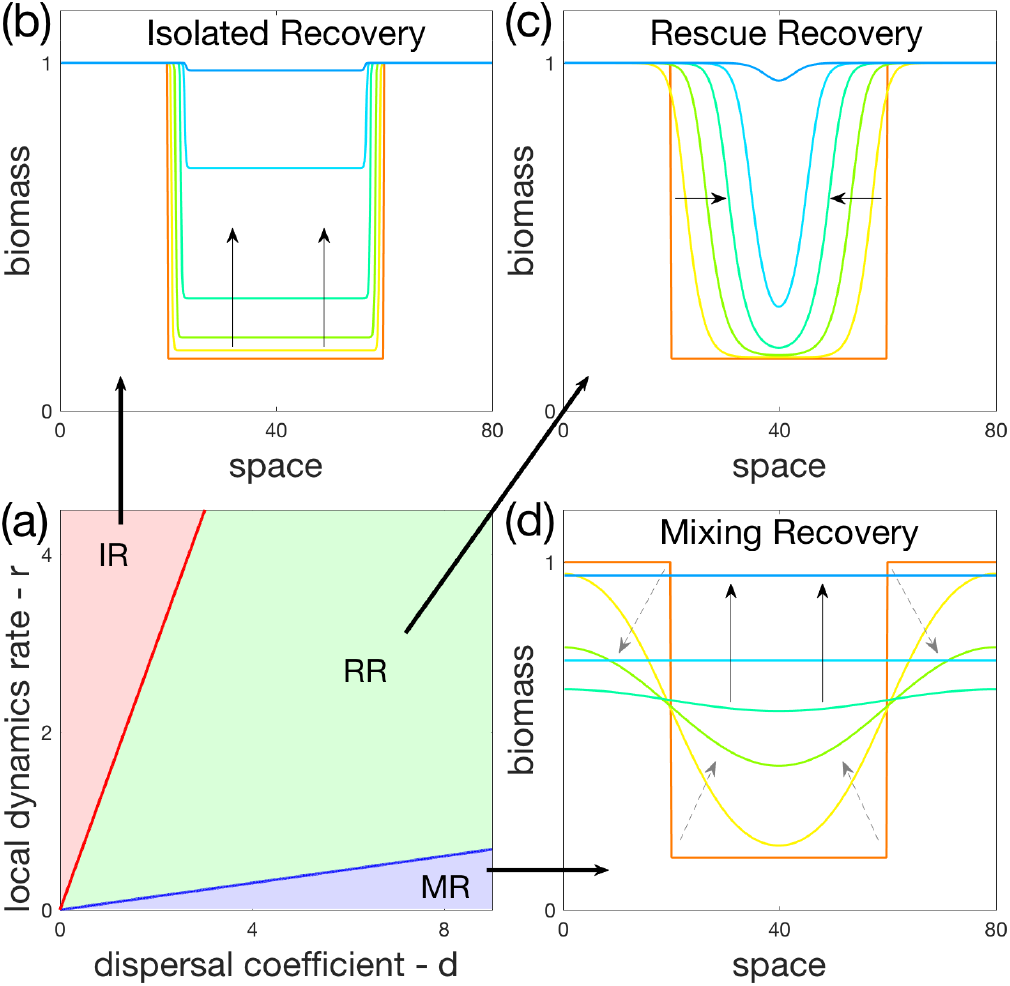
Three recovery regimes across the space of possible systems. **(a)** parameter space of local dynamics rate *r* versus dispersal coefficient *d*, noting the three regimes of recovery: Isolated (IR), Rescue (RR), Mixing (MR). **(b,c,d)** Spatial profiles of biomass at different times (shown by different colors), demonstrating the recovery under each regime, with black arrows showing the direction of recovery. The system’s equilibrium of *N* = 1 is disturbed by setting half the system to *N* = 0.15 (orange), with different colors (yellow to blue) showing the recovery over time. **(b)** IR (*d* = 0.01, *r* = 4) shows recovery due to local processes alone.In RR (*d* = 1, *r* = 1) recovery due to spatial spread dominates. In MR (*d* = 8, *r* = 0.01) the system first homogenizes (gray dashed arrows), and then at a later time the recovery takes place.

**FIG. 2:**
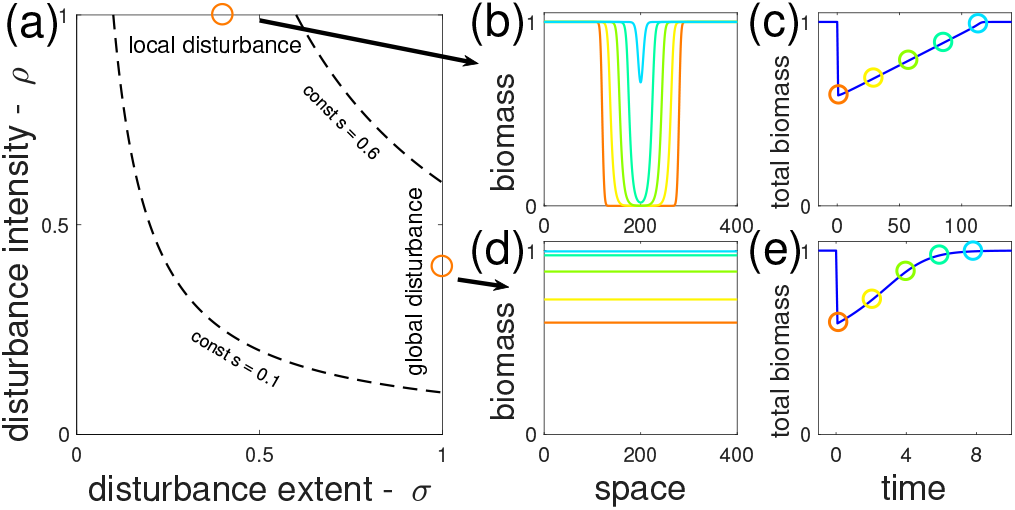
Space of possible disturbances, and the recovery that follows them. **(a)** parameter space of disturbances: spatial extent (*σ*) versus local intensity (*ρ*). The disturbance’s overall strength is *s* = *σρ*, where dashed curves show constant values of s. **(b,c)** Recovery following a local disturbance (*σ* = 0.4, *ρ* = 1). **(d,e)** Recovery following a global disturbance (*σ* = 1, *ρ* = 0.4). Spatial profiles of biomass at different times (orange through blue) are shown in (b,d), while in (c,e) the overall biomass over time is shown (with circles of corresponding colors to the profiles shown in (b,d)).

## III. RECOVERY REGIMES

Knowing that different system and disturbance properties can lead to different recovery behaviors, we now seek a more systematic understanding of the relationship between these properties and recovery. To quantify the recovery process, we study its return time, defined as the duration needed for the system to recover 90% of lost biomass. In Fig. 3 we map out return time as a function of disturbance extent, disturbance intensity, and dispersal. While return time grows with all three disturbance properties, *ρ, σ*, and *s*, there are substantial differences in how they affect return time as we increase dispersal.

When dispersal is weak (Fig. 3a), the effect of neighboring regions is minimal, so that the dynamics are mostly governed by local processes. In particular, if the intensity *ρ* is not too large then local recovery dynamics are fast, so that one can ignore spatial effects, and the return time is effectively set by *ρ*. However, for very high *ρ* there is a switch in behavior, and the spatial extent *σ* becomes dominant in determining return time. This is because local recovery is now slow enough to allow for spread processes to become relevant, and their timescale is largely set by *σ*. For intermediate dispersal (Fig. 3b), the spatial processes can have more of an effect, so that *σ* becomes important at lower values of *ρ*. Hence, we see here the same picture as in Fig. 3a, except that the switch in behavior occurs for lower *ρ*.

Finally, for strong dispersal (Fig. 3c) spatial processes are fast enough to homogenize the system before local dynamics become significant. Therefore the spatial structure of the disturbance is irrelevant, only its overall strength *s* is important.

In fact, we can understand these dependencies on disturbance properties by knowledge of the recovery regimes (note that magenta lines in Fig. 3 show the transition lines between regimes). In IR (low *ρ* for low and intermediate *d*) the recovery is entirely local so that return time is controlled by *ρ*. In RR (high *ρ* for low and intermediate *d*) spatial spread processes are responsible for recovery so that return time is controlled by *σ*. Finally, in MR (high *d*) recovery only takes place after the disturbance has spread throughout the system, hence *s* controls return time.

We now take a broader look at the three recovery regimes, and in particular on the transitions between them. As suggested by the conceptual diagram of Fig. 1, we expect that the dispersal *d* and local dynamics *r* play a similar but opposite role in determining which type of recovery takes place. Thus IR would occur for low *d* (high *r*), MR would occur for high *d* (low *r*), and RR may occur for intermediate values of both. We expect the system size *L* to have a similar role to *r*, since for smaller systems lower dispersal levels would be needed to homogenize the system. This general intuition is validated by measuring the partial contribution of each regime to the recovery, as demonstrated by the parameter space of *d* and *L* in Fig. 4a, see Appendix B.

**FIG. 3:**
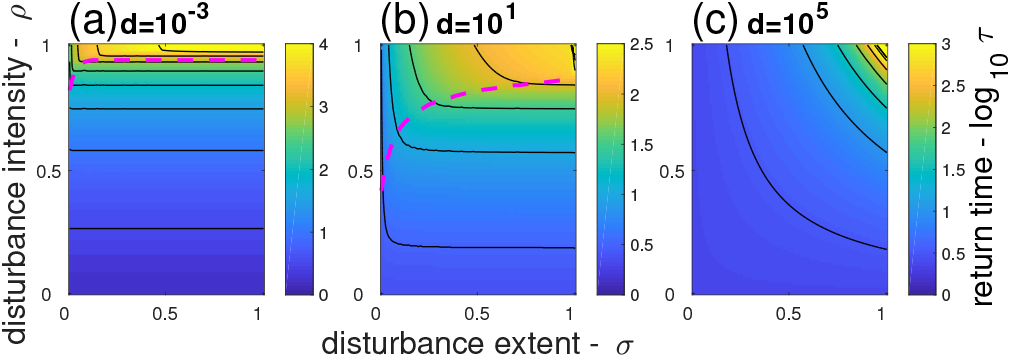
Return time across the disturbance space for three levels of dispersal. Each panel shows return time *log_10_τ* as a function of disturbance extent *σ* and intensity *ρ*. Dispersal increases from left to right as:*d* = {10^-3^,10^1^, 10^5^}. **(a,b)** IR and RR below and above magenta line (low and high *ρ*), respectively; **(c)** MR. Black lines show contours at 0.5 intervals.

**FIG. 4:**
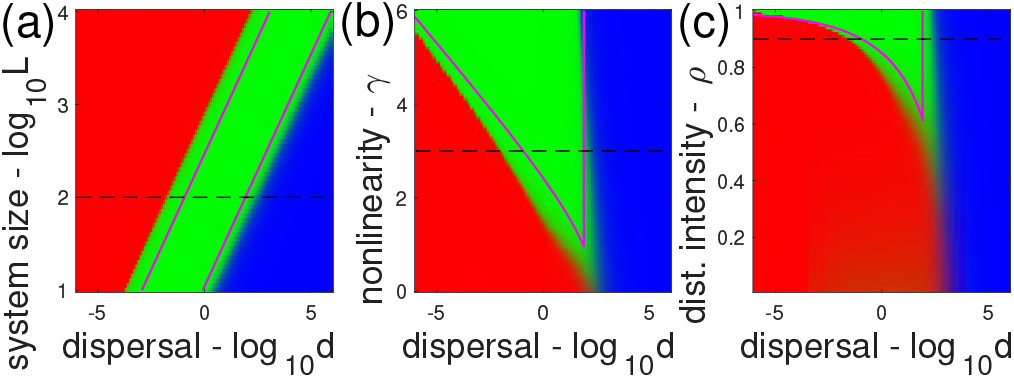
Effects of dispersal versus system size, nonlinearity of local dynamics and disturbance intensity on the three recovery regimes. IR (red), RR (green) and MR (blue) regimes shown on three parameter spaces. The parameter spaces span the dispersal coefficient *d* on the x-axis (in log scale), and the **(a)** system size *L*, **(b)** nonlinearity of local dynamics *γ*, **(c)** intensity of disturbance *ρ*, on the y-axis. Magenta lines show the prediction of the transitions between regimes (see main text for details). Dashed black line in each panel corresponds to the same set of parameters: *L* = 100, *γ* = 3, *ρ* = 0.9.

A final defining feature of the recovery regimes is the ratio between the timespans of local and regional recovery. Since the system may be far from its equilibrium locally, the timespan of local recovery may be very large, even infinitely so, depending on the disturbance intensity *ρ* and the system nonlinearity, typified by *γ* in our model. By increasing either *ρ* or *γ*, local recovery in the disturbed region slows down considerably, while rescue dynamics are essentially unchanged. This is because rescue dynamics are driven by a flow of biomass from an undisturbed region, unaffected by the slow dynamics far from equilibrium. Therefore RR grows at the expense of IR, as seen in Fig. 4b,c.

We now formalize the heuristic reasoning described so far. Consider the dynamics of two adjacent uniform domains, one in equilibrium and the other is a disturbed domain away from the equilibrium. Due to dispersal a smooth transition region (front) will form between these two domains. Such fronts have two main consequences for recovery: their motion into the disturbed domain leads to rescue dynamics, and hence RR, while if the fronts themselves are so large that they take over the entire system, then the system is in MR. If the fronts do neither of these things while recovery takes place, then the system is in IR. As detailed in Appendix A, we use these implications of front dynamics [39] on recovery, combined with a dimensional analysis [22], to reach a prediction of the transitions between the recovery regimes. The result can be summarized by visualizing an axis of effective system size 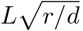. As this effective size grows, at 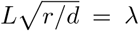 the system goes from MR to RR and at 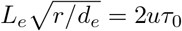 from RR to IR.

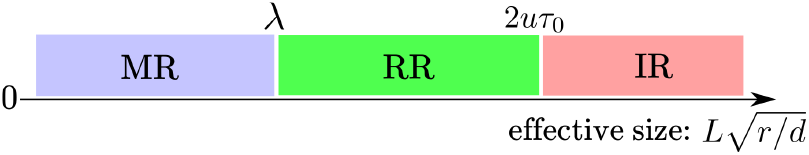

The three physical parameters, *L, r* and *d*, are the same as described previously, while the three other quantities, *u, λ* and *τ*_0_ are nondimensional properties of the model. *u* and λ are the front speed and size, which do not depend strongly on model or disturbance properties (with typical values in the order of 1 and 10 respectively). On the other hand, *τ*_0_ is the recovery time without dispersal (scaled by *r*), with values that can change by orders of magnitude, depending on characteristics of both system (*γ*) and disturbance (*ρ*).

The prediction of regime transitions are shown in magenta lines in Fig. 4. The dimensional parameters *d, r* and *L* all occur in the same term, implying a similar role in determining the recovery regime, as seen by the parallel lines in Fig. 4a due to changes in *d* and *L*. On the other hand, the changes in *τ*_0_ (Fig. 4b,c) due to increasing either *γ* or *ρ* lead to a larger region in parameter space where RR is dominant instead of IR. However, these same changes in *γ* and *ρ* do not effect much the values of *u* and *λ*, so that the RR-MR transition remains largely unchanged.

## IV. EXAMPLES

Our framework of recovery regimes can be applied to a wide range of questions in ecology, such as stability in a fragmented landscape and the interplay of synchrony and local extinctions. We detail here a few concrete examples and discuss their implications, to both demonstrate the application of the framework, and give credence to its generality. In doing this we will also show that our initial assumptions, focusing on dynamics of a single species around a stable equilibrium in one spatial dimension, do not in fact limit the predictive power of the framework.

### A. Metapopulation stability

Two recent experiments [9, 15] studied the consequences of a localized population extinction on a metapopulation, the first by Dai et al. [9] in order to consider a new spatial indicator of impending collapse, and the second by Gilarranz et al. [15] to understand how modularity of the spatial structure can buffer disturbances. Despite apparent similarities in the details of these studies, they are actually relevant in different recovery regimes, and hence in entirely different settings. Dai et al. considered a bistable system, where the dynamics of populations close to the tipping point slow down, leading to larger “recovery length”, a measure of how large is the front that connects a disturbed region from a non-disturbed region. The slowing down of dynamics near the tipping point effectively means that the system has smaller *r*, which would lead it from IR, to RR and on to MR. However, in practice the proposed indicator is limited to RR, since in MR the “recovery length” is larger than the system, so no notable change can be detected, while IR is irrelevant due to the bistability of the system. On the other hand, Gilarranz et al. looked at how decimating a population locally leads to a propagation of the disturbance across the system. In this case the assumption is that the disturbance does not directly propagate (as opposed to the spread of an infection, for example), but that due to spatial movement of the population the other parts of the system still suffer from the disturbance. As can be seen in the different recovery panels in Fig. 1, this behavior only occurs in MR, where the front is of a size that is comparable to the system. Overall, these two examples together show that knowing the recovery regime of a given system is paramount to the applicability of different ecological indicators.

### B. Fragmentation scenarios

A more complex spatial setting is that of habitat fragmentation, which is a leading factor of biodiversity loss [5, 19, 33]. As both the number of habitat patches and the number of potential dispersal routes shrink due to fragmentation, the effects these changes have on various ecosystem properties, such as biodiversity and stability, are of high interest. Taking this question into our framework, we may expect that such changes in spatial structure will change both the effective dispersal in the ecosystem, and its effective size. We could therefore relate different scenarios of fragmentation to trajectories in the parameter space of Fig. 4a, where the dispersal coefficient should be related to the average link number per site *α*, and the system size should be related to the average shortest path between two sites *β* (See Appendix C for details). As shown in Fig. 5a, by starting with a random spatial network, and removing sites from the system, different behaviors emerge. If sites are taken out from the periphery, then both *α* and *β* shrink (Fig. 5a, dashed line), leading to a tight-knit system and therefore to MR (Fig. 5b). On the other hand, if the sites are taken out randomly, then *β* tends to grow (Fig. 5a, solid line), leading to IR (Fig. 5c). Thus, a change in spatial structure can lead to a complete switch in the system’s dynamical behavior.

**FIG. 5:**
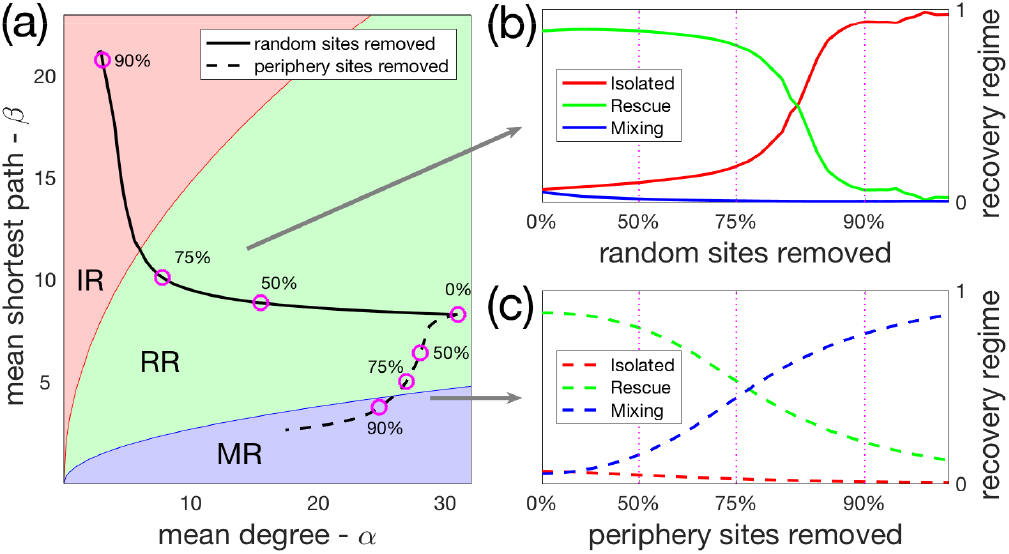
Fragmentation scenarios leading to a change in recovery regimes. **(a)**Parameter space of mean degree *α* (average number of links per site, as a proxy for dispersal coefficient *d*) versus mean shortest path *β* (proxy for system size *L*). Overlaid are two scenarios of fragmentation, one of randomly taking out sites (solid line) and one of taking out periphery sites (dashed line). Background colors are based on a prediction of transition lines between recovery regimes. **(b,c)** Contribution of the three recovery regimes to the overall recovery in a network, along two scenarios of fragmentation. Horizontal axis shows (in a logarithmic scale) the percentage of sites removed either (b) randomly or (c) from periphery. Magenta circles in (a) show the removal of {0%, 50%, 75%, 90%} of sites, with corresponding vertical lines in (b,c). Initial network has 2000 sites, with approximately 40 links per site on average, and: *r* = 1, *d* = 2, *γ* = 3, *ρ* = 0.7, *σ* = 0.5. See more details in Appendix C.

### C. Predator-prey synchronization

A different aspect of stability in a spatial setting, is that of synchrony and how it is affected by dispersal [1, 17, 32]. Extinction due to synchrony of populations is of particular concern when the local dynamics do not have a stable equilibrium, as often seen in predator-prey interactions. Recent work [12] has experimentally shown that inducing localized extinction events can stabilize systems since it prevents spatial synchronization. Such synchronization occurs in MR, since it is here that the dispersal is strong enough to homogenize the system, and in fact the RR-MR transition coincides with the definition of synchronization length [21]. We can therefore expect that localized extinctions will be helpful in a very specific setting, one where synchronization is weak enough so that localized extinctions can prevent it, and that rescue dynamics are strong enough to allow the system to recover from the extinctions themselves. Hence we can predict that disturbances will increase survival around the RR-MR transition, as can indeed be seen in Fig. 6 (see Appendix D for details). We are thus able to pin-point the parameters for which this unintuitive phenomenon will occur. Moreover, this example highlights the fact that exotic behavior can be expected when a system is crossing the boundaries between recovery regimes.

**FIG. 6:**
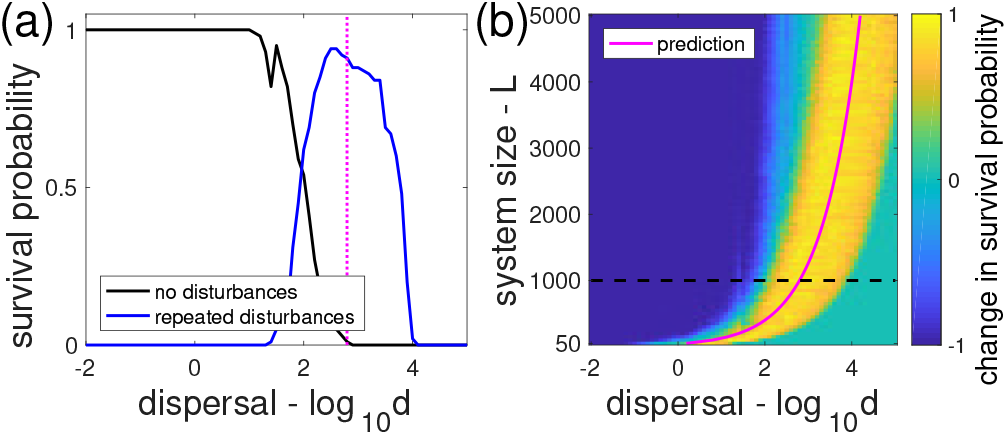
Localized extinction disturbances increase stability due to prevention of synchronization. Following Fox et al. [12], we model populations of predator and prey in a spatial settings, which would go extinct without dispersal. **(a)** Survival probability of predator as a function of dispersal coefficient *d*. Results for no disturbances (repeated disturbances) are shown in black (blue). **(b)** The change in survival probability (by the addition of repeated extinction disturbances) in a parameter space of dispersal coefficient *d* and system size *L*. The RR-MR transition line (magenta) predicts where disturbances increase survival. Black dashed line shows the location of the cut shown in (a), *L* = 1000. See more details in Appendix D.

### D. Metacommunity biomass productivity

Our framework can be applied beyond the context of stability, for example by looking at productivity in a metacommunity context [23]. As shown in recent work on metacommunity dynamics [35], along the dispersal axis one can see three different mechanisms responsible for biomass production, base growth, species sorting and mass effect (Fig. 7a). We can relate these three different mechanisms of biomass production to the three recovery regimes.

**FIG. 7:**
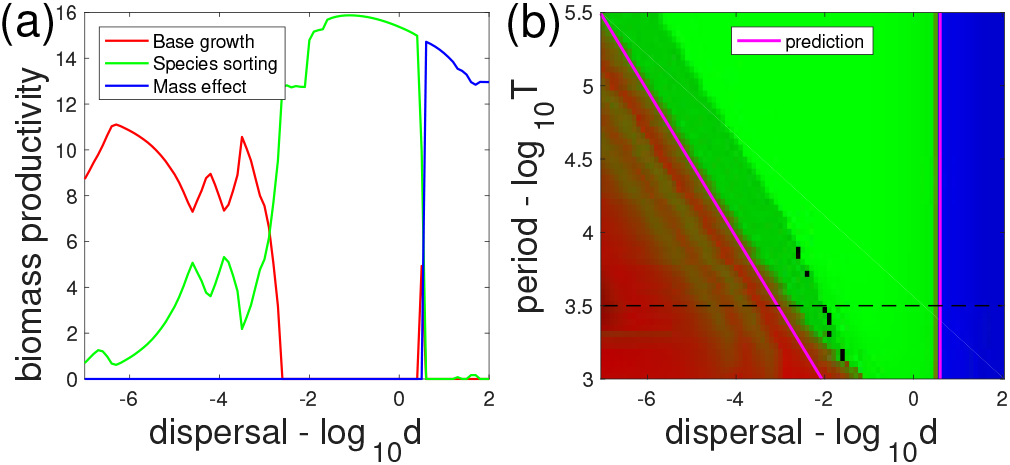
Biomass productivity in a metacommunity due to three different mechanisms. Following Thompson et al. [35], we model an ecosystem of multiple species and a single resource in a spatial settings, with periodically changing conditions. **(a)** Biomass productivity as a function of dispersal coefficient *d*, where the (red, green, blue) lines correspond to three different mechanisms: base growth, species sorting, mass effect. **(b)** Parameter space of dispersal coefficient *d* and period of environmental change *T*, showing the different mechanisms. Magenta lines show the prediction of the transition lines between the three recovery regimes, which correspond nicely to the three mechanisms of biomass productivity. Black dashed line shows the location of the cut shown in (a), *T* = 3200. See more details in Appendix E.

For weak dispersal, biomass production will be due to base growth, where communities are effectively isolated and the species composition does not follow changes in the environmental conditions. Thus base growth corresponds to IR, in which local dynamics (of competitive exclusion and recovery) are faster than spatial spread. This behavior is contrasted with species sorting that occurs with higher dispersal, where species do not occur throughout space at a given time, but rather follow their optimal conditions. Species sorting corresponds to RR, in which spread processes allow species to follow conditions, and the system to recover. Finally, for strong dispersal, species are abundant due to mass effects, by which species biomass from highly productive locations is dispersed across space. Mass effects correspond to MR, in which dispersal is significantly faster than local dynamics, so that both biomass and disturbance spreads throughout the system. Due to this correspondence between mechanisms of biomass production and recovery regimes, we can predict which mechanism will be dominant depending on the system’s parameters, as shown by the magenta curves in Fig. 7b. To predict the transitions we replace recovery time without dispersal with the period of environmental change, since the latter sets the local timescale (see Appendix E for details). This example highlights that basic dynamical processes, such as those captured by our approach, underly the behavior of complex ecological systems. In our case, understanding these processes sheds light beyond recovery properties, predicting changes in ecosystem functioning.

## V. DISCUSSION

By analyzing the response to a disturbance of a simple yet spatially explicit model, we could make powerful predictions about the stability properties of various complex systems, ranging from metacommunities in a changing environment to populations in a fragmented landscape. We have found three regimes of recovery, Isolated (IR), Rescue (RR) and Mixing (MR), and shown how these regimes depend on both the properties of the system, and the disturbance that is imposed on it. We could therefore map these regimes to a space comprised of the parameters of both the system and the disturbance. The processes involved in the recovery from a disturbance in each of these regimes are qualitatively different. For instance, in MR the system first homogenizes before local processes drive the system back to equilibrium, whereas in RR the recovery is driven by the propagation of biomass from undisturbed regions into the disturbed ones. For this reason the relationship between time to recovery and disturbance property (extent, intensity and strength) differs substantially between regimes. More generally, any prediction about the effect of a given parameter on the system’s stability will strongly depend on the regime the system is in.

From a mapping of these regimes we could predict the effects of fragmentation and global change on basic features of ecosystem stability. What matters in this context is not only where the system is on the map, but where it is going. When global change moves the system closer to collapse, local dynamics slow down, and therefore the system is pushed towards MR. This is the premise of various early warning signals of catastrophic transitions (e.g. recovery length of Dai et al.), made explicit within our framework. These indicators essentially measure how close the system is from MR (although being in this regime does not imply a collapse). Our mapping is especially useful when considering the combined effect of fragmentation and global change, since fragmentation may push the system towards IR, in contrast to the effect of slowing down due to global change. This means that the recovery regime may not change at all, so that early warning signals that measure the distance to MR will not detect the impending collapse. A much clearer picture is gained from knowledge of the system’s trajectory on the map of recovery regimes.

This work demonstrates that despite inherent complexities and details in the dynamics of populations and ecosystems, strong qualitative predictions can be made from the analysis of a simple and generic model. Indeed, we studied the dynamics of a single species at equilibrium in a uniform one-dimensional landscape, and considered its response to a single disturbance. However, our example of fragmentation scenarios shows that we can apply our methodology to more complex spatial structure. Moreover, the predictions shown for the predator-prey and the metacommunity systems clearly show that our framework can be applied to multiple species systems. These two examples also show that considering a system disturbed from equilibrium did not limit our predictive ability, as the predator-prey system was one where multiple disturbances occurred and the dynamics exhibited large oscillations, while the metacommunity system had no explicit disturbance at all, but rather a continuously changing environment. Overall these examples suggest that there are universal properties of ecosystem dynamics in a spatial settings, that can be unraveled from dimensional considerations.

To determine the three recovery regimes, we propose that there are two main constants to consider. The first is an effective size, determined by three dimensional parameters of the system {*d,r,L*}, which combine to form a nondimensional constant 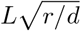. When this constant is very small 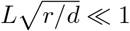, then the system is effectively well-mixed, and therefore in MR. When this is not the case, a measure of endogenous resilience 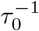 needs to be considered, where *τ_0_* was defined as the relative (e.g. nondimensional) recovery time without dispersal, which depends on local dynamics and disturbance intensity. If the product of both constants is large, then the system acts as multiple isolated sites, and hence is in IR. Otherwise, if endogenous resilience is small while effective size is still large, then the system is in the Recovery regime, in which undisturbed domains of the system are the main instigator of recovery and thus stability.

In general, determining the relevant nondimensional constants should not be expected to be a straightforward task. However, this defines a clear direction towards a deeper understanding of the spatial processes in a system of interest. For example, in our analysis of the fragmentation scenarios the effective system size was essentially the average shortest path between two sites, rather than the total number of sites in the network. The issue of estimating the relevant parameters should be particularly interesting in the context of complex species interactions, due to collective emergent behavior. For instance, the characteristic timescale of a community, such as its local recovery time, may be an emergent property of the assembly process. Moreover, in the case of strong heterogeneities between species (e.g. substantially different dispersal abilities between trophic levels) it may be that more nondimensional constants are needed to faithfully describe the system’s dynamics. For instance, a higher trophic level may spread much faster than its resource [29], leading to a combination of IR and MR.

It is worth noting that only in RR can we expect to see an interplay of spatial scales, between the characteristic scale of the system and any scale that is imposed on it. This leads us to ask what other phenomena we can find in this regime, especially since this regime is not well explored in comparison with IR and MR. In particular, the transition regions between RR and the other two recovery regimes can be expected to be especially interesting and display exotic behavior, since there is no clear timescale separation. Indeed, the surprising results shown by Fox et al., where local extinctions save the predator population from total extinction, well demonstrate this notion that the transition regions can show particular and unintuitive behavior.

This study is essentially based on the presumption that we can learn a great deal on an ecological system by performing a simple calculation using our knowledge of its dimensional properties. The examples we have presented show that this claim has merit, and our methodology can indeed provide new insights into dynamics of spatial systems. This study is an important first step in determining the possible responses of spatially extended ecological systems to disturbances, and draws for the first time a clear link between the opposite settings of a well-mixed system and a set of isolated sites. Considering the proliferation of theoretical and empirical studies, and the growing sets of observational and experimental data, being able to qualitatively compare different systems is of vital importance. This study suggests a simple way of performing such a mapping, using limited information about a given system. Such an approach is relevant for both theoretical models and empirical data, paving the way towards a novel synthetic view on ecosystem dynamics in space.

## Acknowledgements

We wish to thank Bart Haegeman for helpful discussions and review of the manuscript, and Matthieu Barbier for helpful discussions. This work was supported by the TULIP Laboratory of Excellence (ANR-10-LABX-41) and by the BIOSTASES Advanced Grant, funded by the European Research Council under the European Union’s Horizon 2020 research and innovation programme (grant agreement No 666971).

## Authorship

YRZ, JFA and ML designed the study, YRZ and JFA performed the research; YRZ, JFA and ML wrote the manuscript.

## Methods

Detailed explanations on the simulations used to create the different figures can be found in Appendices B through E.

All simulations used to create the figures were made using Matlab with the library RDM: https://github.com/yzelnik/RDM

Code that shows how simulations and calculations for each figure were made can be found at: https://github.com/yzelnik/trsr-scripts

## Appendix A: Dimensional analysis of front dynamics

We describe here the analysis preformed on the model given by equation 1, which will lead us to the predictions of recovery regimes. To do this we use insight on the dynamics of fronts, combined with a dimensional analysis, as described below.

We begin by assuming that following a disturbance the system is composed of two domains, one at equilibrium, and a disturbed domain that is away and potentially very far from this equilibrium. Between these two domains smooth transitions regions (fronts) will form that will connect the two domains, due to dispersal. These fronts have two main consequences on recovery: their motion into the disturbed domain leads to rescue dynamics, and hence RR, while if the fronts themselves are so large they take over the entire system, then the system is in MR. If the fronts do neither of these things while recovery takes place, then the system is in IR. We can therefore predict that the transition between IR and RR occurs when the timespan of local recovery *T_0_* is the timespan of regional recovery *T_R_*. The latter can be approximated by the ratio between the system size *L* and twice the front speed *U* (since two fronts take part in the recovery), leading to: *L*/(*2U*) = *T_R_*. The transition between RR and MR occurs when the front size Λ is as large as the system, and hence *L* = Λ.

We now rewrite equation 1 by considering one spatial dimension, and defining new non-dimensional variables for space, time and biomass {*x, t, n*} instead of the old dimensional variables {*X, T, N*}, to produce equation A1:

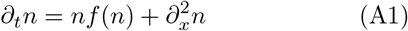

Where the definition of the new variables are: *x* = 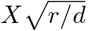; *t* = *Tr*; *n* = *N/K*, and we rewrite the *F* function as *f(n)* = *F*(*nK*). Since equation A1 has the same structure as equation 1, they share the same properties, and in particular the same general dynamics of fronts. We therefore note the non-dimensional front speed and size as *u* and λ respectively. Due to dimensional constraints, we know that the relations between the new and old front properties are: 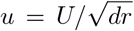 and 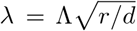 (note that front properties do not depend on the system size *L*). Both *u* and λ are non-dimensional constants, and thus unrelated to any dimensional properties such as *d, r* or *L*. They therefore depend only on the type of local dynamics, as described by *f(n)*, and moreover, can be expected to be of the order of 1 for all different models. We can also use this definition of *u* and λ, together with a non-dimensional constant of the recovery time without dispersal *τ_0_* = *T_0_r* to redefine the boundaries between the three recovery regimes:

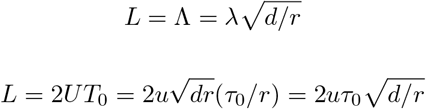

We therefore have two inequalities for the transitions between the three regimes:

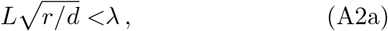

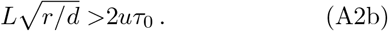

where the prediction is that if the first inequality holds then the system is in MR, if the second inequality holds then the system is in IR, and otherwise the system is in RR. Note that we implicitly assume that both inequalities cannot hold, namely that

## Appendix B: Analysis of simple model

The results shown in Fig. 1–4 are all based on equation B1, in one spatial dimension.

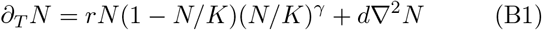

In general, there are seven parameters relevant for these results. The five parameters of the system are: local dynamics rate *r*, dispersal coefficient *d*, carrying capacity *K*, nonlinearity *γ*, and system size *L*. Two parameters define the disturbance: its local intensity *ρ*, and its spatial extent *σ*, where the overall strength of the disturbance is *s* = *σρ*.

For Fig. 1, in the three simulations shown the parameters *r* and *d* were varied to showcase the three different recovery regimes. (b) IR was shown using *r* = 4, *d* = 0.01, (c) RR was shown using *r* = 1, *d* = 1, and (d) MR was shown using *r* = 0.01,d = 8. Other parameters were held constant: *K* = 1, *γ* = 3, *L* = 80, *ρ* = 0.85, *σ* = 0.5.

In Fig. 2 two simulations were used, where (b,c) the localized disturbance had *σ* = 0.4, *ρ* = 1, while (d,e) the global disturbance had *σ* = 1, *ρ* = 0.4, so that the overall strength of the disturbance was the same with *s* = 0.4. Other parameters were held constant: *r* = 1, *d* = 1, *K* = 1, *γ* = 3, *L* = 400.

For Fig. 3 the three parameter spaces shown were calculated with different levels of dispersal: (a) *d* = 10^-3^ (b) *d* = 10^1^ (c) *d* = 10^5^. In all, the extent *σ* and intensity *ρ* were varied between 0 and 1. Other parameters were held constant: *r* = 0.5, *K* = 1, *γ* = 3, *L* = 500.

For Fig. 4, in all three parameter spaces shown the dispersal coefficient *d* was varied (in log scale) between 10^-6^ and 10^-6^. For panel (a) the system size *L* was varied (in log scale) between 10 and 10000, while *γ* = 3, *ρ* = 0.9. For panel (b) the nonlinearity parameter *γ* was varied between 0 and 6, while *L* = 100, *ρ* = 0.9. For panel (c) the disturbance intensity *ρ* was varied between 0. 01 and 1, while *L* = 100, *γ* = 3.

For both Fig. 1 and Fig. 4 the transition lines of IR-RR and RR-MR are shown, based on the analytical formula 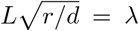 and 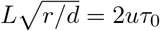, respectively. To do this three non-dimensional constants were estimated numerically, using simulations of the non-dimensional equation A1 (effectively setting: *r* = 1, *d* = 1, *K* = 1). By simulating a front between *n* =1 and *n* = 0, the front speed *u* was estimated as *u* = 0.37 and the front size was estimated as λ = 11. Note that here we always used *γ* = 3 and *ρ* = 1 (in effect, since one domain had *n* = 0), since the dependence of both λ and *u* on these changes in minimal. On the other hand, for estimation of τ_0_ both *γ* and *ρ* were very significant. Here we looked at a comparable system, but we set the dispersal to zero *d* = 0, and estimated the recovery time (to just below *N* =1) when the entire system is set to a value 1 — ρ. For *γ* = 3, *ρ* = 0.85 (as used in Fig. 1) we had τ_0_ = 132, while for *γ* = 3, *ρ* = 0.9 (as used in Fig. 4a) we had τ_0_ = 400.

## Appendix C: Fragmentation scenarios

For the analysis of fragmentation scenarios, we used the same general modeling scheme of equation B1, as detailed in the previous section, but instead of a onedimensional system, a spatial network was used. In this network setup, we used link-wise dispersal, so that each link allows for a proportional amount of biomass to disperse, and hence the more links a site has, the more biomass will disperse between it and its neighbors. Each link is identical, and so is each site in its local conditions. Dispersal level was defined so that if a network is made of a one dimensional chain, and the number of sites in the network *M* is the same as the size of a one dimensional system *L*, then the dispersal (and in general all other results) will be identical between these two systems (assuming dispersal is not too strong). This was done simply for an easier comparison with results of a one dimensional system, and has no bearings on the results since it amounts to a scaling of the dispersal coefficient. In practice, the dynamical equation per site is:

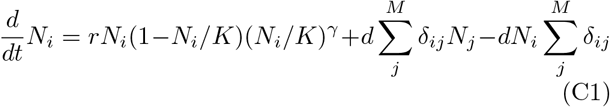

where *N_i_* is the biomass at site *i*, and *δ_ji_* is the connectivity matrix, with values of 1 if there is a link between sites *i* and *j*, and 0 otherwise.

The networks used were constructed in the following way. *M* sites were given random locations (uniform distribution) in a two-dimensional space within a circle. For each site, the *c* closest neighboring sites were chosen, and if a link did not already exist, then it was formed between the initial and neighboring site with a probability *p*. In all simulations the parameters used were *M* = 2000, *c* = 40, *p* = 0.5, so that the average number of links per site was approximately 31.

To simulate different fragmentation scenarios, two methods of removing sites were used. The scenario where periphery sites were removed (solid lines in Fig. 5) was done by looking at the distance of each site from the center of the whole network. Here the locations are defined as the ones originally used to construct the network. The sites that were farthest from the center were removed, in a non-random fashion. The second scenario was that of removing random sites (dashed lines in Fig. 5). Here an iterative scheme was used to avoid a disconnection of the network to several clusters due to the site removal. In this way the network remains one cluster so that biomass can propagate from each site to any other site. This was done so that the effective size of the network could be assessed faithfully using the measure of the average shortest path (see below).

The iterative scheme is as follows. At each iteration step, *M* sites are chosen at random. If their removal will not disconnect the network, then the removal is enacted. If it will disconnect the network, then no sites are removed in this step, and instead the value of *M* is halved, as long as it is not already smaller than a threshold value of *z*. The iterations are repeated until the required number of sites are removed, or up to a maximum number of iterations *i*. If this maximum is reached (not enough sites were removed) than the remaining number of sites to be removed are chosen at random, ignoring the issue of network disconnection. For a network site of *M* = 2000 sites, we started with *M* that is the total number of sites to be removed, with a threshold value *z* of 0.002 of the overall number of sites to be removed, and maximum iteration number *i* = 500. Using this iterative scheme, the network does not disconnect until over 90% of sites are removed, whereas a completely random removal would disconnect the network at around 70% — 80% of sites removed. At the same time, other properties of the network, such as its recovery dynamics, do not change considerably.

The two axes shown in Fig. 5a are the average link number in the x-axis and the average shortest path in the y-axis. The average link number is a proxy for the dispersal coefficient of the system *d*. It is calculated as the number of links a given site has, averaged over all sites in the network. The average shortest path is a proxy for the size of the system *S*. It is measured by calculating the minimal number of sites that are needed to connect between two given sites, averaged over all possible pairs of two sites. If a site is not connected, so that some sites are not reachable by others, than the average shortest path is defined as infinity. In Fig. 5a such networks are not shown.

The simulations of the fragmentation scenarios were preformed similarly to those of the one-dimensional system, except that equation C1 was used, with *δ_ij_* which are the result of a network creation and site-removal processes described above. The disturbance was imposed by choosing an initial site at random, and going through its neighbors and their neighbors and so forth, until *M_σ_* sites were chosen, such that *M_σ_* = *σM*. We used *σ* = 0.5, so that *M_σ_* = 1000 sites were affected by a disturbance. Other parameters used were *r* = 1, *d* = 2, *K* = 1, *γ* = 3, *ρ* = 0.7. The results shown were averaged over 100 different networks that were created and underwent the process of site-removal, with a different randomization key. For each network, 10 recovery processes were measured, each with a different initial site for the disturbance.

The transitions between different regimes, as shown in Fig. 5a, are based on the prediction detailed in the previous section. This result has to be augmented to fit the settings of network fragmentation. First, a relation needs to be drawn between the effective dispersal coefficient *d_e_*, which is used instead of *d*, and the average number of links 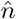. Since the number of links in a chain topology (which is equivalent to a 1D setting given in the previous section) is 2 links per site, we need to normalize by this value, so that 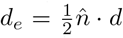. Second, for the effective system size L_e_, we use the average shortest path 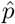 as a proxy for system size *L*. Since the networks we use are set in a two-dimensional (2D) circular space, and the average distance between two points in a such a 2D space is approximately a quarter of the diameter of such a region, we need to normalize by this value. Hence we use: 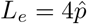. Finally, the homogenizing effect of the front (relevant for the RR-MR transition) needs to be normalized since we are dealing with a 2D space (as opposed to the 1D setting in the previous section). The proper normalization for the front size (λ_e_) is not trivial, and we leave it for future work. However, using geometric considerations we can hypothesize that the normalization constant will be π, and this value indeed gives a good agreement with a mapping of recovery regimes on multiple networks with different parameters that we preformed (not shown). We therefore use λ_e_ = πλ. Putting all these together we have our prediction for the two transition lines: 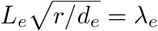 and 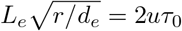

## Appendix D: Predator-Prey system

The analysis of the effect of local disturbances on the stability of a predator-prey metacommunity was done by following previous work on such a system that used both experiments and simulations [12]. We use the third model described in this study, a version of the Rosenzweig-MacArthur predator-prey model. The model used is:

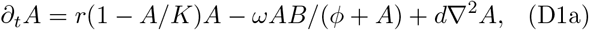

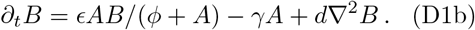

where *A* is the prey biomass and *B* is the predator biomass, and the parameters values used are (following the original paper): *r* = 1.85, *K* = 35920, *ω* = 25.5, *ϕ* = 11364, *∈* = 12.4, *γ* = 2.07 and *d* is varied (see below). The model was simulated in one spatial dimension of length *L* with periodic boundary conditions.

To allow extinctions we check the biomass density at each location every time-step, and if it is smaller than a threshold value of *N_th_* = 1 than its value is set to 0 (biomass densities typically go over 10^3^, so that this has a minimal effect when populations levels are high). In each simulation, the initial population of both predator and prey was randomly chosen from a uniform distribution between 0 and 10^4^ without correlations in space. The simulation was run for a constant length of time *T* = 100, and the predator population was deemed to have survived if it is non-zero anywhere in the system. The simulations were repeated 100 times with different randomization keys, and the results of survival probability derived from averaging over these simulation results.

Two simulation conditions were compared, one where no local disturbances were induced, and the other where 100 disturbances occurred throughout the simulation time in constant intervals, a disturbance every 1 unit of time. This comparison was done while changing two system parameters: *d* was varied (in log scale) between 10^-2^ and 10^5^ and *L* was changed between 50 and 5000.

The prediction shown in Fig 6 of the RR-MR transition uses an estimation of the front size. This was done by taking a system without any predator or prey population, and seeding a small domain of the system with both predator and prey (with density *A* = *B* = 1). While the dynamics in the original domain tend to be chaotic, both prey and predator slowly invade the bare-region. Thus a front is formed after a sufficient time which has the same properties regardless of initial conditions, and its size can be calculated. Since the front in this system is non monotonic and includes spatial oscillations, its estimation is not very accurate, but its size can still be approximated, and a value of 40 was used to produce the transition line shown in Fig. 6 (magenta curve).

## Appendix E: Biomass productivity in a metacommunity

The analysis of biomass productivity in a metacommunity was done by following previous work on such a system [35]. The model used is based on older work of the spatial insurance hypothesis [25], and uses a system with *M* species *N_i_*, that differ in their use of a single resource *R*. The model equations are:

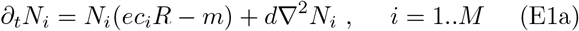

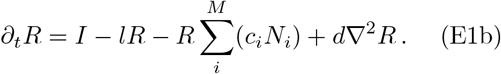

where *e* = 0.2, *m* = 0.2, *I* = 150, *l* = 10, *M* = 9, *L* = 100, dispersal coefficient *d* is varied between simulations (in log scale) in the range 10^-7^ and 10^-2^, and the consumption rates *c_i_* are varied during the simulation itself to simulate temporal and spatial changes in the environmental conditions. This change in environmental conditions occurs with a period of *T_c_* which is varied (in log scale) between 10^3^ and 10^5.5^. Each species has a different periodic curve of its consumption as a function of time and space, so that by either moving through the whole system (distance *L*) or waiting a time period of *T_c_*, the species will experience all possible conditions specified by this curve. More explicitly:

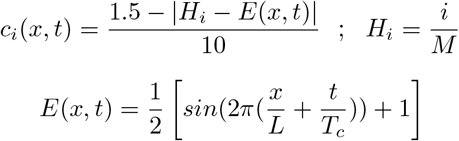

Note that the environmental conditions change with a speed of *v* = *L/T_c_*

For each point in the parameter space of *d* and *T_c_* (Fig. 7), one simulation was made. The initial conditions used were that of uniform densities of all species and resource with an arbitrary value of 1. The simulation was run for a time of 10*T_c_* to minimize the effect of transients, and then an additional time of 5*T_c_* was run, over which the biomass productivity was analyzed (see below). In order to allow species to go extinct, throughout the simulation, at every times-step the biomass of each species is checked at every location, and if it lower than a threshold value of *N_th_* = 0.1 its value is set to 0.

Biomass productivity was defined as the change in biomass at a given time (that is, the right hand side of equation E1a), except for the contribution of mortality (ignoring the term –*mN_i_*), integrated over some region (either the entire system, or some part of it). The biomass productivity was divided between three sources: base-growth, species-sorting and mass-effect. If a species had non-zero densities at every location of space throughout the entire simulation (after the transient run-time of 10*T_c_* was finished), then all its biomass growth was considered to be due to mass-effect. If a species in a certain location had gone extinct (*N_i_* = 0) at any time point in the simulation (but was non-zero in some other time) then all the biomass of this species that was produced throughout the simulation was considered to be due to species-sorting. Productivity in any other case (namely, if a species is always present at a given location, but is not present throughout the entire system) was considered to be due to base-growth.

The prediction shown in Fig 7 of the IR-RR and RR-MR transitions (magenta curves) uses three dimensionless constants: front size, front speed and recovery time. In the context of the this system, there is no explicit disturbance, but rather the constant change of environmental conditions effectively acts as a disturbance. Therefore, instead of a recovery time we use as a proxy the period over which the environmental conditions change, so that τ_0_ = *T_c_*. To estimate the front properties we consider the same system but with a constant environment (*c* = 0.1), and put in one species in a small domain, and follow its propagation into the bare region. From this simulation we can estimate the front properties as: λ = 40 and *u* = 0.54.

## References

[1] Abbott, K. C. (2011), ‘A dispersal-induced paradox: Synchrony and stability in stochastic metapopulations’, Ecology Letters 14(11), 1158–1169.

[2] Arnoldi, J.-F., Bideault, A., Loreau, M. & Haegeman, B. (2018), ‘How ecosystems recover from pulse perturbations: A theory of short-to long-term responses’, Journal of Theoretical Biology 436, 79–92.

[3] Arnoldi, J.-F., Loreau, M. & Haegeman, B. (2016), ‘Resilience, reactivity and variability: A mathematical comparison of ecological stability measures’, Journal of Theoretical Biology 389, 47–59.

[4] Baskett, M. L., Weitz, J. S. & Levin, S. A. (2007), ‘The Evolution of Dispersal in Reserve Networks’, The American Naturalist 170(1), 59–78.

[5] Bennie, J., Hodgson, J. A., Lawson, C. R., Holloway, C. T., Roy, D. B., Brereton, T., Thomas, C. D. & Wilson, R. J. (2013), ‘Range expansion through fragmented landscapes under a variable climate’, Ecology Letters 16(7), 921–929.

[6] Bjørnstad, O. N., Ims, R. A. & Lambin, X. (1999), ‘Spatial population dynamics: Analyzing patterns and processes of population synchrony’, Trends in Ecology & Evolution 14(11), 427–432.

[7] Burkey, T. V. (1989), ‘Extinction in Nature Reserves: The Effect of Fragmentation and the Importance of Migration between Reserve Fragments’, Oikos 55(1), 75.

[8] Connor, E. F. & McCoy, E. D. (1979), ‘The statistics and biology of the species-area relationship’, The American Naturalist 113(6), 791–833.

[9] Dai, L., Korolev, K. S. & Gore, J. (2013), ‘Slower recovery in space before collapse of connected populations’, Nature 496(7445), 355–358.

[10] Donohue, I., Hillebrand, H., Montoya, J. M., Petchey, O. L., Pimm, S. L., Fowler, M. S., Healy, K., Jackson, A. L., Lurgi, M., McClean, D., O’Connor, N. E., O’Gorman, E. J. & Yang, Q. (2016), ‘Navigating the complexity of ecological stability’, Ecology Letters 19(9), 1172–1185.

[11] Fischer, J. & Lindenmayer, D. B. (2007), ‘Landscape modification and habitat fragmentation: A synthesis’, Global ecology and biogeography 16(3), 265–280.

[12] Fox, J. W., Vasseur, D., Cotroneo, M., Guan, L. & Simon, F. (2017), ‘Population extinctions can increase metapopulation persistence’, Nature Ecology & Evolution 1(9), 1271–1278.

[13] Franks, P. J. (1997), ‘Spatial patterns in dense algal blooms’, Limnology and Oceanography 42(5part2), 12971305.

[14] Galiana, N., Lurgi, M., Claramunt-López, B., Fortin, M.-J., Leroux, S., Cazelles, K., Gravel, D. & Montoya, J. M. (2018), ‘The spatial scaling of species interaction networks’, Nature Ecology & Evolution p. 1.

[15] Gilarranz, L. J., Rayfield, B., Linan-Cembrano, G., Bascompte, J. & Gonzalez, A. (2017), ‘Effects of network modularity on the spread of perturbation impact in experimental metapopulations’, Science 357(6347), 199201.

[16] Gouhier, T. C. & Guichard, F. (2007), ‘Local disturbance cycles and the maintenance of heterogeneity across scales in marine metapopulations’, Ecology 88(3), 647–657.

[17] Gouhier, T. C., Guichard, F. & Gonzalez, A. (2010), ‘Synchrony and stability of food webs in metacommunities’, The American Naturalist 175(2), E16–E34.

[18] Haegeman, B. & Loreau, M. (2014), ‘General relationships between consumer dispersal, resource dispersal and metacommunity diversity’, Ecology Letters 17(2), 175184.

[19] Hanski, I. & Ovaskainen, O. (2000), ‘The metapopulation capacity of a fragmented landscape’, Nature 404(6779), 755–758.

[20] Kerr, B., Riley, M. A., Feldman, M. W. & Bohannan, B. J. (2002), ‘Local dispersal promotes biodiversity in a real-life game of rock-paper-scissors’, Nature 418(6894), 171.

[21] Lande, R., Engen, S. & Sæther, B.-E. (1999), ‘Spatial Scale of Population Synchrony: Environmental Correlation versus Dispersal and Density Regulation’, The American Naturalist 154(3), 271–281.

[22] Legendre, P., Loic F. J. Legendre, TotalBoox & TBX (2012), Numerical Ecology., Elsevier Science. OCLC: 969016657.

[23] Leibold, M. A., Holyoak, M., Mouquet, N., Amarasekare, P., Chase, J. M., Hoopes, M. F., Holt, R. D., Shurin, J. B., Law, R., Tilman, D., Loreau, M. & Gonzalez, A. (2004), ‘The metacommunity concept: A framework for multi-scale community ecology: The metacommunity concept’, Ecology Letters 7(7), 601–613.

[24] Loreau, M. & de Mazancourt, C. (2013), ‘Biodiversity and ecosystem stability: A synthesis of underlying mechanisms’, Ecology Letters 16, 106–115.

[25] Loreau, M., Mouquet, N. & Gonzalez, A. (2003), ‘Biodiversity as spatial insurance in heterogeneous landscapes’, Proceedings of the National Academy of Sciences 100(22), 12765–12770.

[26] MacArthur, R. (1955), ‘Fluctuations of Animal Populations and a Measure of Community Stability’, Ecology 36(3), 533.

[27] May, R. M. (1973), Stability and Complexity in Model Ecosystems, number 6 *in* ‘Monographs in population biology’, Princeton University Press, Princeton, N.J.

[28] McCann, K. S. (2000), ‘The diversity-stability debate’, Nature 405(6783), 228–233.

[29] McCann, K. S., Rasmussen, J. B. & Umbanhowar, J. (2005), ‘The dynamics of spatially coupled food webs: Spatially coupled food webs’, Ecology Letters 8(5), 513523.

[30] Plitzko, S. J. & Drossel, B. (2015), ‘The effect of dispersal between patches on the stability of large trophic food webs’, Theoretical Ecology 8(2), 233–244.

[31] Qian, H., Badgley, C. & Fox, D. L. (2009), ‘The latitudinal gradient of beta diversity in relation to climate and topography for mammals in North America’, Global Ecology and Biogeography 18(1), 111–122.

[32] Ripa, J. (2000), ‘Analysing the Moran effect and dispersal: Their significance and interaction in synchronous population dynamics’, Oikos 89(1), 175–187.

[33] Saunders, D. A., Hobbs, R. J. & Margules, C. R. (1991), ‘Biological Consequences of Ecosystem Fragmentation: A Review’, Conservation Biology 5(1), 18–32.

[34] Scheffer, M. & Carpenter, S. R. (2003), ‘Catastrophic regime shifts in ecosystems: Linking theory to observation’, Trends in Ecology & Evolution 18(12), 648–656.

[35] Thompson, P. L., Rayfield, B. & Gonzalez, A. (2017), ‘Loss of habitat and connectivity erodes species diversity, ecosystem functioning, and stability in metacommunity networks’, Ecography 40(1), 98–108.

[36] Turner, M. G. (1989), ‘Landscape Ecology: The Effect of Pattern on Process’, Annual Review of Ecology and, Systematics 20(1), 171–197.

[37] von Hardenberg, J., Kletter, A. Y., Yizhaq, H., Nathan, J. & Meron, E. (2010), ‘Periodic versus scale-free patterns in dryland vegetation’, Proceedings of the Royal Society B: Biological Sciences 277(1688), 1771–1776.

[38] Wang, S., Loreau, M., Arnoldi, J.-F., Fang, J., Rahman, K. A., Tao, S. & de Mazancourt, C. (2017), ‘An invariability-area relationship sheds new light on the spatial scaling of ecological stability’, Nature Communications 8, 15211.

[39] Zelnik, Y. R. & Meron, E. (2018), ‘Regime shifts by front dynamics’, Ecological Indicators.

